# The Role of CD147 in Leukocyte Aggregation in Liver Injury

**DOI:** 10.1101/600676

**Authors:** Christine Yee, Nathan Main, Alexandra Terry, Igor Stevanovski, Annette Maczurek, Alison J. Morgan, Sarah Calabro, Alison J. Potter, Tina L. Iemma, David G. Bowen, Golo Ahlenstiel, Fiona J. Warner, Geoffrey W. McCaughan, Susan V. McLennan, Nicholas A. Shackel

## Abstract

**Background:** Chronic inflammation is the driver of liver injury resulting in progressive fibrosis and eventual cirrhosis. The consequences include both liver failure and liver cancer. We have previously described increased expression of the highly multifunctional glycoprotein CD147 in liver injury. This work describes a novel role of CD147 in liver inflammation and the importance of leukocyte aggregates in determining the extent of liver injury.

**Methods:** Non-diseased, progressive injury and cirrhotic liver from humans and mice were examined using mAb targeting CD147. Inflammatory cell subsets were assessed by multicolor flow cytometry.

**Results:** In liver injury, we observe abundant intrahepatic leukocyte clusters defined as ≥5 adjacent CD45^+^ cells which we have labelled “leukocyte aggregates”. We have shown that these leukocyte aggregates are significant in determining the extent of liver injury. If CD147 is blocked *in vivo,* these leukocyte aggregates diminish in size and number together with a marked significant reduction in liver injury including fibrosis. This accompanied by no change in overall intrahepatic leukocyte numbers. Further, blocking aggregation formation occurs prior to an appreciable increase in inflammatory markers or fibrosis. Additionally, there were no observed, “off-target” or unpredicted effects in targeting CD147.

**Conclusion:** CD147 mediates leukocyte aggregation which is associated with the development of liver injury. This is not a secondary effect, but a cause of injury as aggregate formation proceeds other markers of injury. Leukocyte aggregation has been previously described in inflammation dating back over many decades but till now been shown to determine the extent of injury.

## Introduction

The classical hallmark of liver injury is the deposition of abnormal/fibrotic extracellular matrix (ECM) and the eventual development of cirrhosis which is mediated by the activated hepatic stellate cell (HSC)^(1)^. Chronic inflammation drives ongoing HSC activation and fibrosis^(1)^. Chronic liver inflammation can be regarded as commencing with an initial innate immune response to an ongoing insult that develops into chronic injury with both sustained innate and adaptive immune components^(2–6)^. T-cell responses are clearly pivotal to the development of chronic immune-mediated hepatic injury and it has now been shown that B-cells are essential for the development of intrahepatic fibrosis leading to cirrhosis^(7–11)^.

Importantly, we have previously identified a number of novel pathways of liver injury^(12–14)^. Arising from these studies we have shown that hepatocytes remodel extracellular matrix in liver injury via production of active matrix-metalloproteinases^(15)^.

CD147 is an abundant 269aa type 1 integral glycosylated multifunctional membrane protein with differing cellular functions on differing cell subpopulations. The concept of multifunctional proteins is well established^(16–20)^ with at least 3% of all proteins in the human protein interaction network being ‘extreme multifunctional’ proteins^(20)^. CD147 is expressed within a wide range of tissues, including endothelium and epithelium^(21–23)^. CD147 is thought to act via MAPK p38, ERK-1, -2, PI3K and NF-KB signaling pathways^(24–27)^. The regulation of CD147 expression is largely uncharacterised.

A striking and consistent function attributed to CD147 is its ability to regulate leukocyte chemotaxis. Cyclophilins (CyP)-A and B are the two CD147 ligands known to mediate chemotaxis^(21, 28–30)^. CD147 has been shown to be important for leukocyte recruitment in rheumatoid arthritis, multiple sclerosis and inflammatory lung disease, with mAb αCD147 interventions leading to reduced neutrophil, T-cell and monocytes/macrophage infiltration^(11, 28, 31–37)^.

CD147 is mostly found in membrane protein complexes^(29, 30, 38–40)^ and many of these binding partners are matrix components or inflammatory mediators that are dramatically increased with injury (i.e. hyaluronan^(41)^, intercellular adhesion molecule (ICAM)-1^(38, 42, 43)^, lymphocyte function-associated antigen (LFA)-1^(44–46)^ and CD43^(47, 48)^). Additionally, CD147 is also found in large complicated multiple protein “super-complexes” on the cell surface containing many binding partners^(39)^. Studies have shown that the domain of CD147 that mediates cell migration is distinct from the domain mediating MMP induction (within extracellular loop I)^(49, 50)^. Therefore, it is well established that CD147 is a multifunctional protein^(16–20)^ with distinct binding partners and structural domains determining its varied functions at specific cellular locations.

Leukocyte aggregations of either lymphoid or myeloid origin are thought to exacerbate inflammation and in turn, the severity of disease. In acute viral infection, TLR stimulation and TNF signaling induced myeloid cell aggregation in the liver^(51)^. These myeloid aggregates induced localized CD8 T cell proliferation. Myeloid cell aggregation was not observed in chronic viral infection. It is yet to be determined if immune cell aggregation occurred in non-viral induced liver injury. The molecules associated with immune aggregation are also unknown. Aggregation of leukocytes within the liver tissue during disease has been most observed in HCV and ALD, though their role in pathogenesis and mode of development is still unclear^(52)^. We have previously shown that CD147 is increased in cirrhotic liver and expressed by hepatocytes (not HSC) during progressive liver injury^(15)^. Herein we report CD147 upregulation on multiple immune cells during liver injury. This manuscript is the first description of a novel CD147 mediated mechanism of immune cell aggregation in liver injury. Specifically, we have shown that CD147 mediated immune cell aggregation is independent of cell proliferation and anti-CD147 intervention reduces immune cell aggregation.

## Materials and Methods

### Human ethics

Human tissues samples were obtained from Royal Prince Alfred Hospital, Sydney with approval of Human Research Ethics Committee (X10-0072). Human tissue used in this study was previously utilized for research^(12, 15)^. Informed written consent was obtained from all participants. The ethics committee waived the need for written consent for use of donor tissue. In Australia, the ethics of human research is governed by the National Statement on Ethical Conduct in Human Research (2007) issued by the National Health and Medical Research Council (NHMRC). Under these guidelines all research involving humans requires ethical approval. Non-diseased donor and end-stage cirrhotic liver tissues were collected from patients attending Prince Alfred Hospital, Sydney during liver transplantation.

### Mice

Mice wild type (balb/c and C57Bl/6) were housed at the Centenary Institute Animal facility in compliance with the Animal Care and Ethics Committee at the University of Sydney (K75/10-2008/3/4801). Liver injury was induced by either thioacetamide (TAA) (300mg/L) (*MP Biomedicals*, Ohio, USA) in drinking water *ad. libitum* for four, eight or twenty weeks or by carbon tetrachloride (CCl_4_) (*Univar, Ajax Chemicals*, Sydney) intraperitoneal injection with 100μl of a 12% CCl4 in paraffin oil mixture twice weekly for four weeks. Control mice were injected with paraffin oil alone. Both treated and age-matched control mice were sacrificed by CO_2_ asphyxiation at conclusion of treatment. CD147 antibody was administered by i.p injection twice weekly (100μg) and control mice received IgG2a (100μg) (HB-189, ATCC). The rat anti-mouse IgG2a CD147 blocking antibody (mAb RL73.2) was produced by hybridoma cells and purified as described^(53)^.

### Gene expression analysis

Total RNA from liver leukocytes, hepatocytes and whole liver were isolated with TRIzol reagent (Invitrogen, San Diego, CA) and cDNA synthesized with SuperScript™ III Reverse Transcriptase (Invitrogen). Quantitative RT-PCR was performed using SYBR® green fluorescent dye (Invitrogen). Specific Taqman probes were used for amplification of CD147 (forward 5’-GTCCAGGAAGTCAACTCCAA-3’; reverse, 5’-GCTCAGGAAGGAAGATGCAG-3’) and this was normalised to housekeeping control 18S (forward, 5’-CGGCTACCACATCCAAGGA-3’; reverse, 5’-CTGGAATTACCGCGGCTG-3).

### Liver function tests

Blood was collected from the inferior vena cava into MiniCollect Serum Tubes [Grenier Bio-One]. Serum was isolated by centrifugation at 6000 *r*evolutions *p*er *m*inute (rpm) for 10 min and the supernatant was collected. Serum was diluted 1:3 in PBS and tested for the activity of enzymes *al*kaline *p*hosphatase (ALP), *a*lanine *t*ransaminase (ALT) and *a*spartate *t*ransaminase (AST) by the Sydney South West Pathology Service. All results are measured in international units per litre (U/L).

### Flow cytometry of CD147 surface expression on leukocyte subsets

Flow cytometry was performed on liver and spleen leukocytes from control and TAA-treated mice. Cells were co-stained with antibodies against CD3ε – PE/Cy7 (145-2C11, BioLegend), CD4-Pacific orange (RM4-5, Invitrogen), CD8α – Pacific blue (53-6.7, BioLegend), CD19 – AlexaFlour 488 (6D5, BioLegend), NK1.1-PE (PK136, Becton Dickinson), F4/80-PerCP/Cy5.5 (RM8, BioLegend) and CD147 – biotin (OX114, BioLegend) further stained with Streptavidin-APC (Invitrogen). Flow cytometry data was collected with FACS LSR-II (Becton Dickinson). Each leukocyte subset was analysed for expression of CD147 and data recorded as median fluorescence intensity. Bar graphs represent mean ± SEM. Statistical analysis was performed by Kruskal-Wallis test followed by Dunn’s multiple comparisons test.

### Immunofluorescence analysis

Indirect immunofluorescence was performed on frozen fixed (acetone:methanol) liver tissue using monoclonal antibody CD45-FITC (1:100) (30-F11, BD Pharmingen). Co-localisation was assessed using CD147 (1:100) (RL73.2, hybridoma), Gr-1 (RB6-8C5, BD Pharminogen) and F4/80 (CI:A3-1, hybridoma), B220 (RA3-6B2, BD Phaminogen), CD19 (ID3, BD Pharmingen) and CD3e (UCHT1, Dako). Secondary antibodies goat anti-rat AlexaFluor® 594 (1:200) (#A-11007, Molecular Probes), goat anti-rabbit AlexaFluor®633 (#A-21071, Invitrogen) were used. For control of background staining, the primary antibodies were omitted or replaced by IgG isotype control (Cat ID: 559073, Dako). The fluorescent images were imaged using a Leica TCS SP5 II confocal microscope (Leica, Georgia, USA).

### Cell cluster quantitation

Liver sections stained for CD45 by immunofluorescence were viewed using Leica DM RBE Fluorescence Microscope [Leica Microsystems] and photographed using Leica DFC 500 camera [Leica Microsystems] in random fields of view (FOV) (n=3-10). Fluorescent cells were quantified blind in using Image-J [National Institute of Mental Health, MD, USA]. Cell clusters were defined as greater than five cells in contact.

### Statistics

Statistical analysis was performed using Prism 6. Mann-Whitney U t-test was performed to compare against control, unless otherwise specified. Significance was accepted at p<0.05. Data is shown as mean ± SEM. As indicated some data is expressed as fold change from control.

## Results

### CD147 gene expression increases in the liver during injury

We have previously observed higher CD147 expression in primary biliary cholangitis (PBC), primary sclerosing cholangitis (PSC) and hepatitis *C* virus (HCV)-induced liver injury in patients ^(12, 54)^. We chose to investigate the expression of CD147 in a TAA-induced mouse model of liver injury to further establish the source and function of CD147 in liver injury. During the development of TAA-induced liver injury we observed a progressive increase in the expression of CD147 mRNA in whole liver tissue (Fig 1A). When hepatocytes and leukocytes were analysed separately a significant increase of CD147 mRNA expression was observed in leukocytes (30-fold) (Fig 1B). Minimal CD147 upregulation was observed on hepatocytes (Fig 1B). This data shows that CD147 expression is upregulated in a mouse model of liver injury and CD147 predominantly increased on leukocytes, not hepatocytes following injury. Further, we examined leukocyte CD147 expression in blood, liver and spleen showing a shift from control to 20 weeks in circulating and intrahepatic leukocytes (Fig 1C).

**Figure 1.**
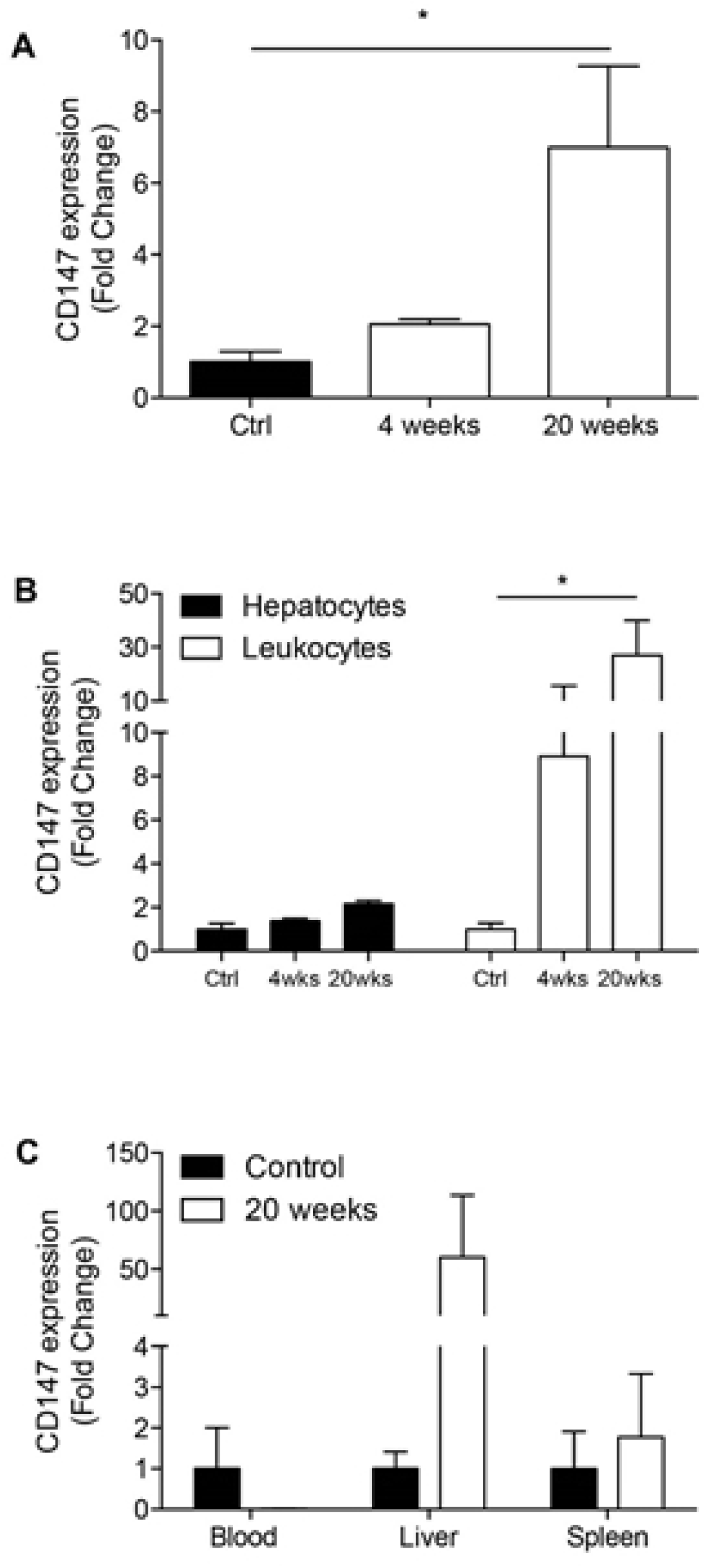
Quantitative Real-Time PCR for CD147 expression in a mouse TAA induced liver injury model. (A) CD147 expression in whole liver from control, 4 week and 20 week TAA-treated mice. (B) Comparison of CD147 expression in liver leukocytes and hepatocytes throughout the course of TAA induced injury. (C) CD147 expression in blood, liver, and spleen at 20 weeks compared to control. Data normalised to 18s, expressed as fold change from control. Bars represent mean+SEM (n=3-6/group). Mann-Whitney test was performed to assess significance from control where * p<0.05.

### CD147 surface expression increases on liver leukocyte subsets during injury

We next characterized the expression of CD147 on various leukocyte subsets during the course of liver injury (Fig 2). Eight colour flow cytometry was utilized to gate CD4^+^ T cells, CD8^+^ T cells, B cells, NK cells, NKT cells, eosinophils, neutrophils and macrophages from the liver. In untreated controls, leukocytes expressed minimal CD147 except for macrophages, neutrophils and eosinophils, which had high basal levels of CD147. CD147 expression increased on all liver leukocytes, with peak expression at 8 weeks of injury. Interestingly, as liver injury progressed, we observed a significant loss of NKT cells and macrophages and a significant increase of CD8^+^ T cells, eosinophils and neutrophils from the liver.

**Figure 2.**
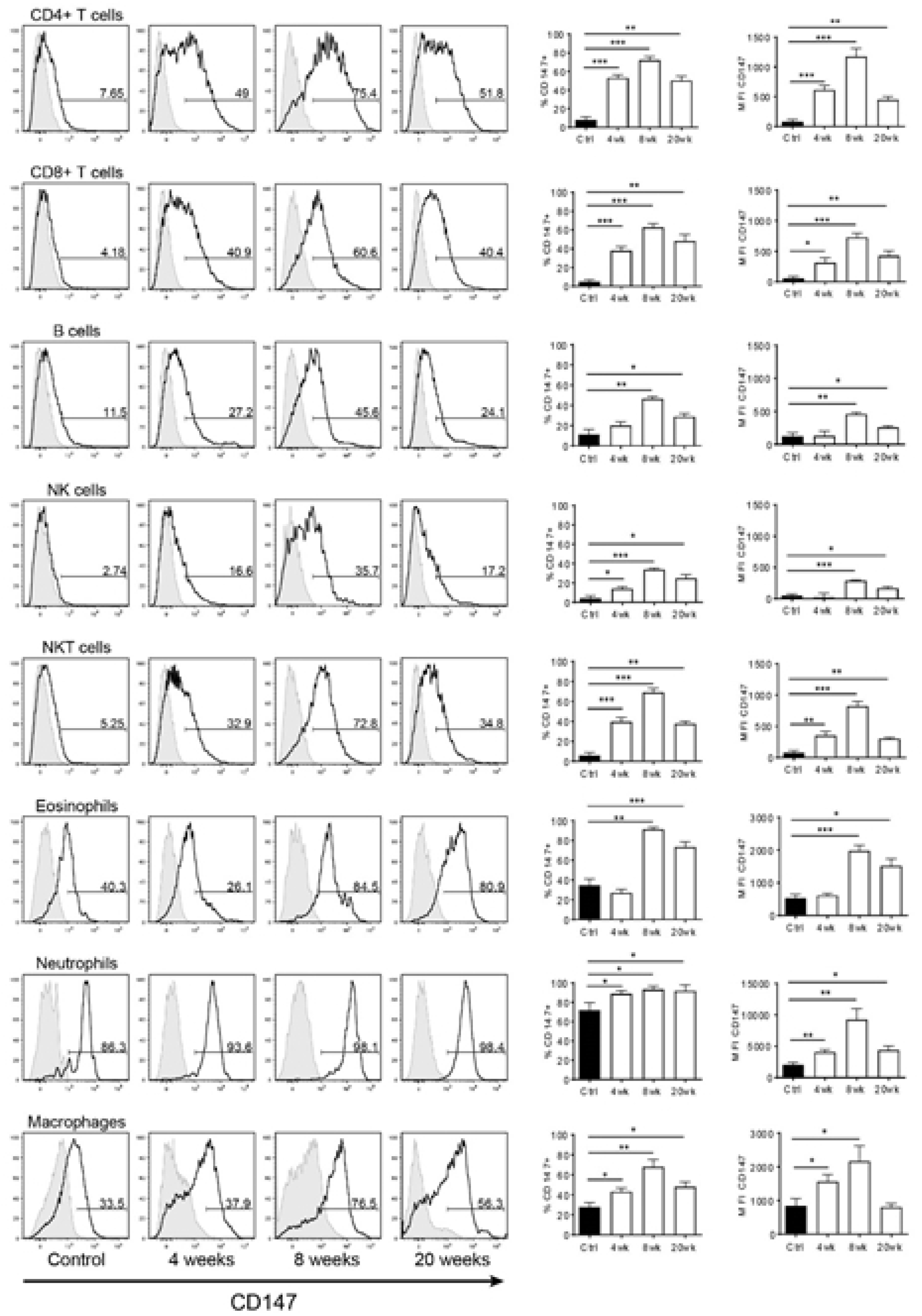
CD147 surface expression on leukocyte subsets isolated from livers in progressive liver injury. Flow cytometry was performed on liver leukocytes from control and TAA-treated mice (n=5-7 per group). Each leukocyte subset was analysed for expression of CD147 and data recorded as percentage of cells positive for CD147 (%CD147+) or mean fluorescence intensity (MFI). Representative histogram plots are shown for each subset, where the black histogram represents the indicated liver subset and grey histogram indicates isotype control. Bar graphs represent mean ± SEM. Mann-Whitney test was performed to assess significance from control where * p<0.05, **p<0.01, ***p<0.001.

To establish whether similar CD147 expression patterns were occurring in an extrahepatic lymphoid organ, the spleen was also analysed by flow cytometry in the same manner described above. This trend was true for all subpopulations examined, with the exception of NK cells, which showed equally high expression at both 8 and 20 weeks. Interestingly, some spleen subpopulations showed a similar trend, however CD147 expression was not nearly as high as in the liver, and no values were significantly up-regulated compared to untreated mice. Within each treatment group, CD147 expression levels were relatively similar amongst all leukocyte subsets, with the exception of macrophages, which displayed expression levels up to 6-fold higher than any other cell type. B cells and NK cells showed the lowest relative expression of CD147, displaying levels half that of the T cell subsets (Fig 2).

### Aggregation of lymphocytes during liver injury in mouse models

To determine how lymphocytes localized during chronic liver injury, CD45 staining was conducted on a range of injury models including TAA treatment in C57Bl/6, CCl_4_ in C57Bl/6 and CCl_4_ in balb/c mice (Fig 3). In uninjured control livers, lymphocytes dispersed evenly within the liver tissue (Fig 3A). In contrast, in both TAA (Fig 3B) and CCl_4_ (Fig 3C) injury models not only were total CD45 cells increased (Fig 3D) but CD45 cells were found to aggregate (defined as >5 CD45^+^ cells in contact). Importantly the majority of aggregation occurred around portal triads where fibrosis first appears. Peak infiltrate was observed after 8 weeks of TAA and 4 weeks of CCl_4_, which correlated with peak immune cell aggregation (Fig 3E). Thus, injury is shown to induce increased immune infiltrate and increased immune cell aggregation.

**Figure 3.**
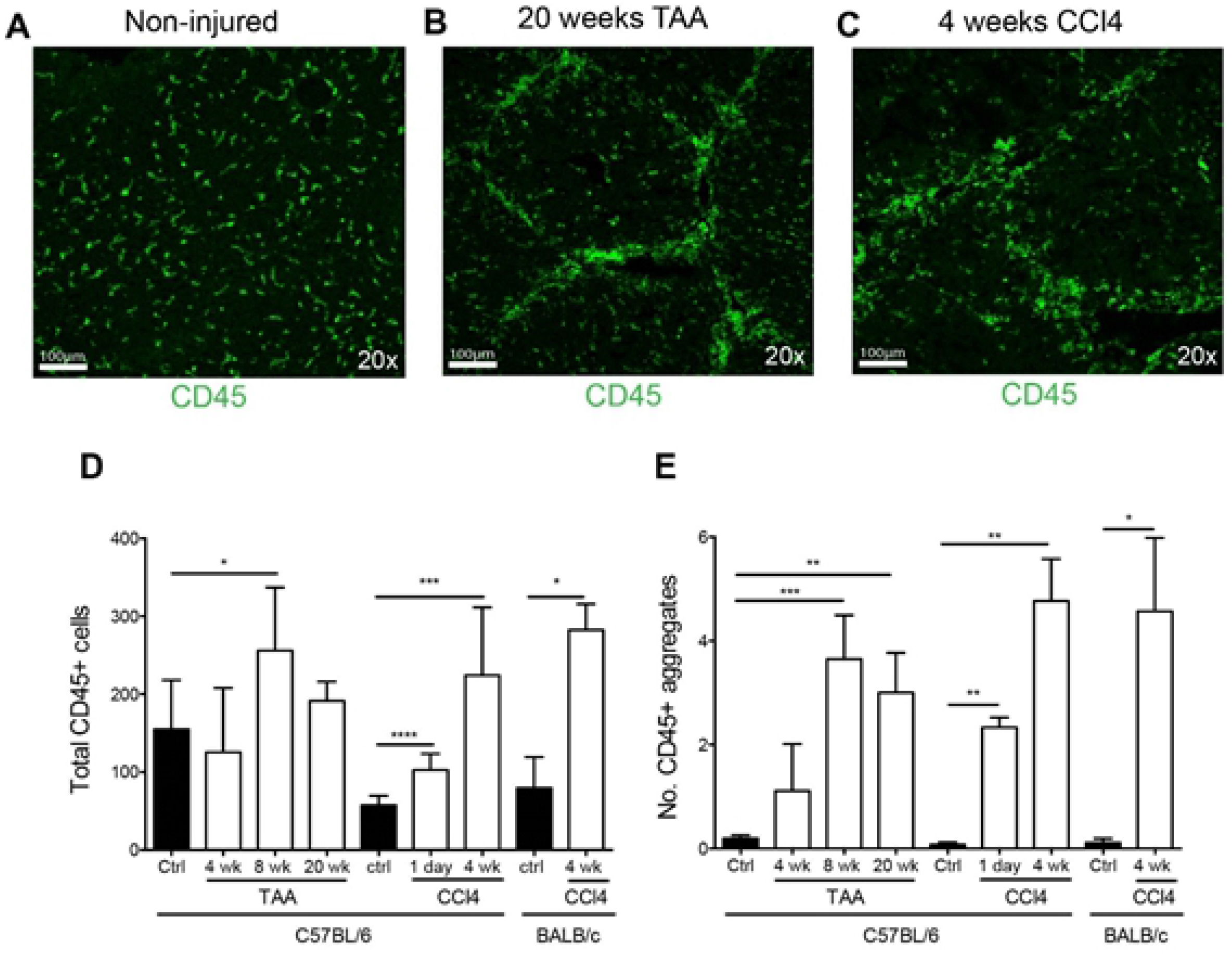
Aggregation of immune cell infiltrate in mouse liver injury. Inflammatory infiltrate in liver tissue of control, 4, 8 and 20 week TAA-treated C56Bl/6 mice and control, 1 day and 4 week CCl_4_ treated C57BL/6 and balb/c mice. Representative images of CD45 stained liver sections from (A) noninjured control; (B) 20 week TAA and (C) 4 week CCl_4_ treated mice. Scale represents 100μM. (D) Total CD45+ cells were quantified per field of view, with 4-10 fields counted per section. Bars represent mean+SEM. (E) Immune cell aggregates were quantified (≥5 CD45+ cells per cluster) per field of view (FOV). Bars represent mean+SEM. Mann-Whitney test was performed to assess significance from control where * p<0.05, **p<0.01, ***p<0.001, ****p<0.0001.

Apoptosis and proliferation within aggregates was assessed by cleaved caspase-3 and Ki-67, respectively (Supplemental Figure 1). This was assessed in both normal and injured liver tissue. In C57Bl/6 mice after 4 weeks of CCl_4_ administration compared to untreated animals, apoptosis went from being most undetected to occasionally seen, which was a significant increase (p = 0.05), but no significant change in proliferation was seen (p = 0. 08). This is consistent with other publications^(55, 56)^.

### Aggregation of lymphocytes in injured human liver

To confirm that immune cell expression of CD147 and immune cell aggregation was not exclusive to mouse models we examined human liver tissue from healthy donors (Fig 4A) and patients with PSC, alcohol induced liver damage (EtOH), AIH and HCV (Fig 4B). Interestingly we not only observed CD147 expression on hepatocytes but all CD45^+^ immune cells expressed CD147 in both healthy and injured livers. Total CD45+ immune cells were significantly increased in diseased liver, irrespective of whether disease was immune mediated (AIH, PSC, HCV) or not (EtOH) (Fig 4C). Inflammatory cells were shown to aggregate with liver injury (Fig 4B) and significantly more immune aggregates were observed in diseased livers compared to healthy donor livers (Fig 4D). In healthy donors although CD45^+^ cells expressed some level of CD147, immune cells did not aggregate. Therefore, this data confirms that in diseased human liver CD147 is expressed on CD45+ immune cell infiltrate in human liver disease and immune cells aggregate during disease.

**Figure 4.**
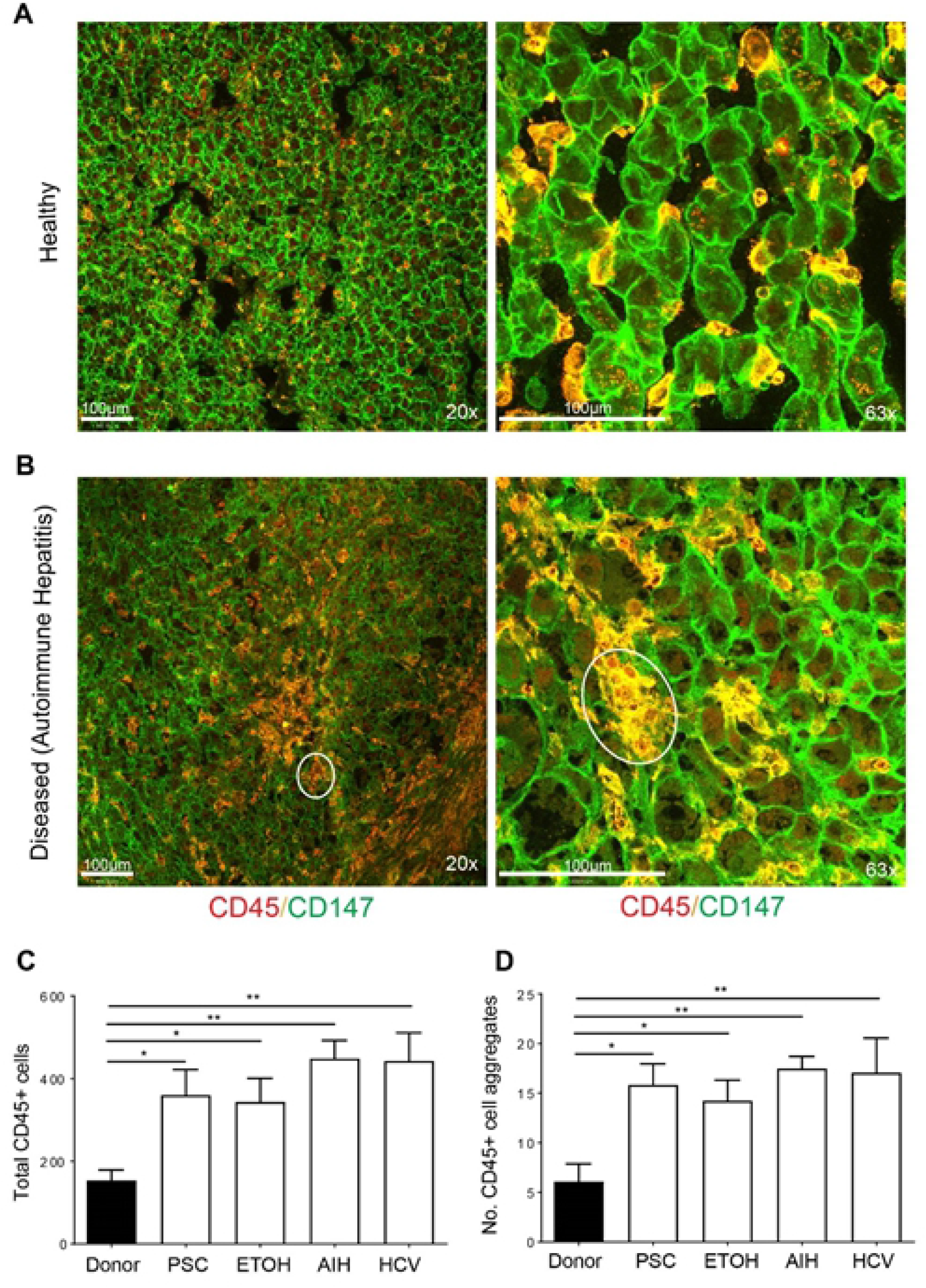
Aggregation of CD147+ immune cells in diseased liver. Representative images of liver sections stained with CD45 (red) and CD147 (green) from (A) healthy control and (B) diseased patient liver (Autoimmune Hepatitis) on either a 20x (left panels) or 63x (right panels) objective. Yellow indicates co-expression of CD147 and CD45. Scale bar represents 100μM. Quantification of (C) total CD45+ cells and (D) immune aggregates in healthy, PSC, EtOH, AIH and HCV livers. Bars represent mean+SEM. Unpaired t-test was performed to assess significance from control where * p<0.05 and **p<0.01. Donor = healthy control liver, PSC = primary sclerosing cholangitis, ETOH = alcohol induced liver damage, AIH = Autoimmune hepatitis, HCV = hepatitis C virus liver injury.

### CD147 dependent leukocyte aggregation in liver injury

To determine if CD147 was important for the formation of immune cell aggregates the effects of anti-CD147 intervention were studied in CCl4 induced liver injury in both C57Bl/6 and balb/c mice. Anti-CD147 did not significantly reduce the number of CD45^+^ cells in the liver (Fig 5E, F). The percentage of immune cells that formed immune cell aggregation was reduced with anti-CD147 intervention in both mouse backgrounds (Fig 5G, H). In C57Bl6 there was non-significant, less than 1.5-fold, increase in ALT with injury and no significant change was seen with anti-CD147 intervention (Fig 5I). However, in balb/c mice with more significant injury the reduction in immune cell aggregation correlated with as significantly reduced serum ALT levels (Fig 5J). Thus, CD147 inhibition appeared to significantly reduce the formation of immune cell aggregates and reduce significant liver injury.

**Figure 5.**
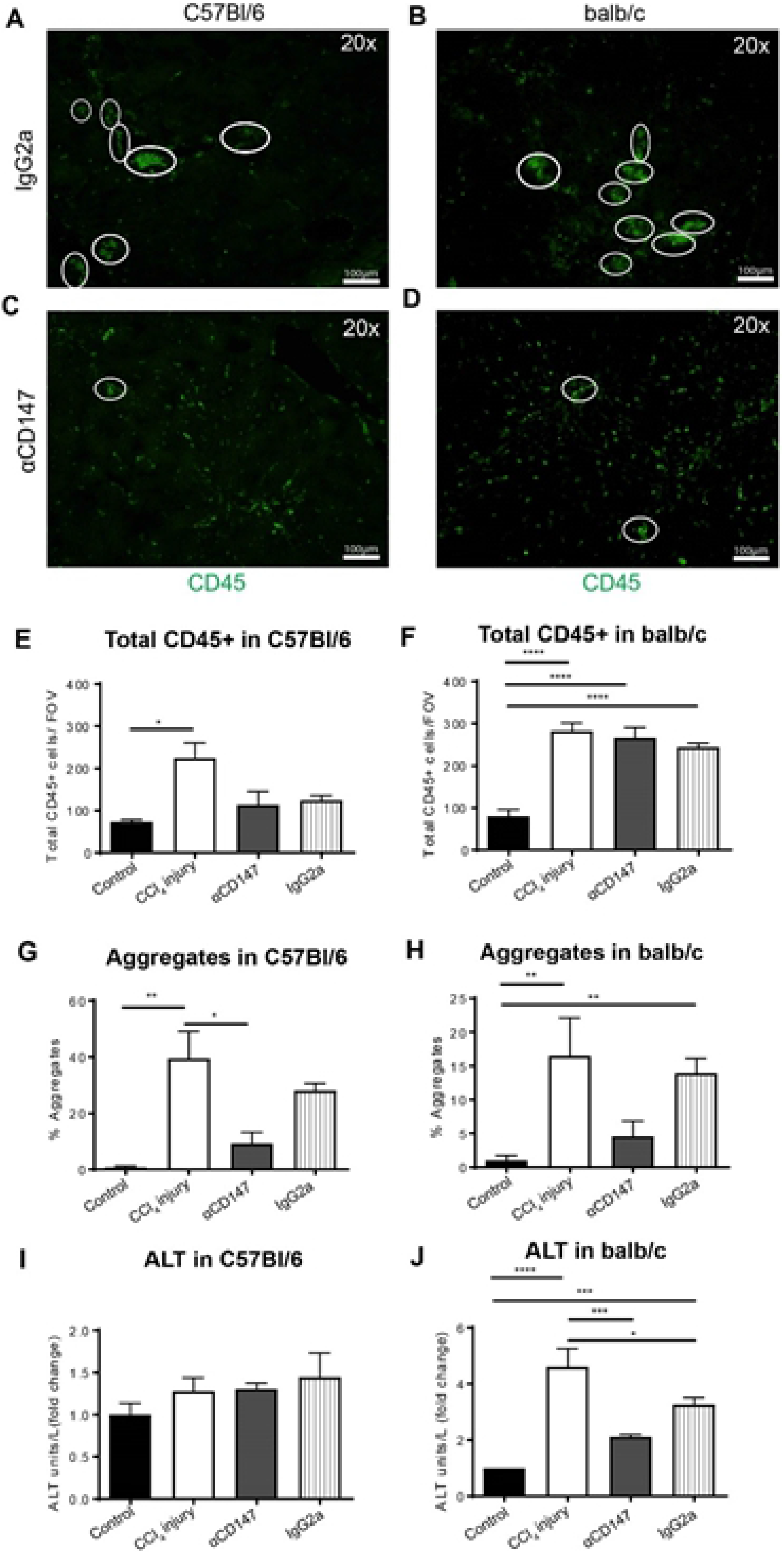
CD147 blockade prevents immune cell aggregation in liver injury. Anti-CD147 antibody intervention in CCl_4_ induced liver injury in C57Bl/6 (left panels) or balb/c (right panels) mice. CD45+ leukocytes were stained in frozen tissue sections in control, treatment (CCl_4_), treatment with αCD147 antibody intervention or treatment with IgG2a isotype antibody controls. Representative immunofluorescent images of CD45+ clusters defined as > 5 cells per cluster (white circles) are shown in C57Bl/6 (A) and balb/c mice (B) treated with IgG2a. Representative immunofluorescence images of CD45+ clusters following αCD147 intervention in C57Bl/6 (C) and balb/c (D) mice. Total CD45+ leukocytes in C57Bl/6 (E) and balb/c (F) mice. (G) Percentage of CD45+ cells in aggregates for each treatment group in C57Bl/6 (G) and balb/c (H) mice. ALT levels (U/mL) as a measure of liver damage in C57Bl/6 (I) and blab/c (J) mice, calculated as a fold change from control. Bar graphs represent mean ± SEM. Mann-Whitney test was performed to assess significance from control where * p<0.05, **p<0.01, ***p<0.001. Control = untreated, CCl_4_ injury = injury alone, αCD147 = anti-CD147 mAb in CCl_4_ injury, IgG2a = Isotype mAb control in CCl_4_ injury.

### Specific leukocyte subsets within aggregates

We have just shown that anti-CD147 intervention reduces the percentage of immune cells in aggregates. To determine what inflammatory cells were aggregating in CCl_4_ induced liver damage and to examine if this is altered during anti-CD147 intervention, liver samples were immunostained with CD45 and F4/80, B220, Gr1 and CD3. F4/80+ Macrophages, B220+ B cells, Gr1+ granulocytes and CD3+ T cells (including NKT cells) were all found in aggregates during liver injury (Fig 6). After anti-CD147 treatment, we observed a decrease in the number of aggregates with at least one F4/80+, B220+ and CD3+ cell. The number of Gr-1+ cells in aggregates was not altered after anti-CD147 treatment (Fig 6). Thus, anti-CD147 intervention was shown to target specific cells reducing immune cell aggregation.

**Figure 6.**
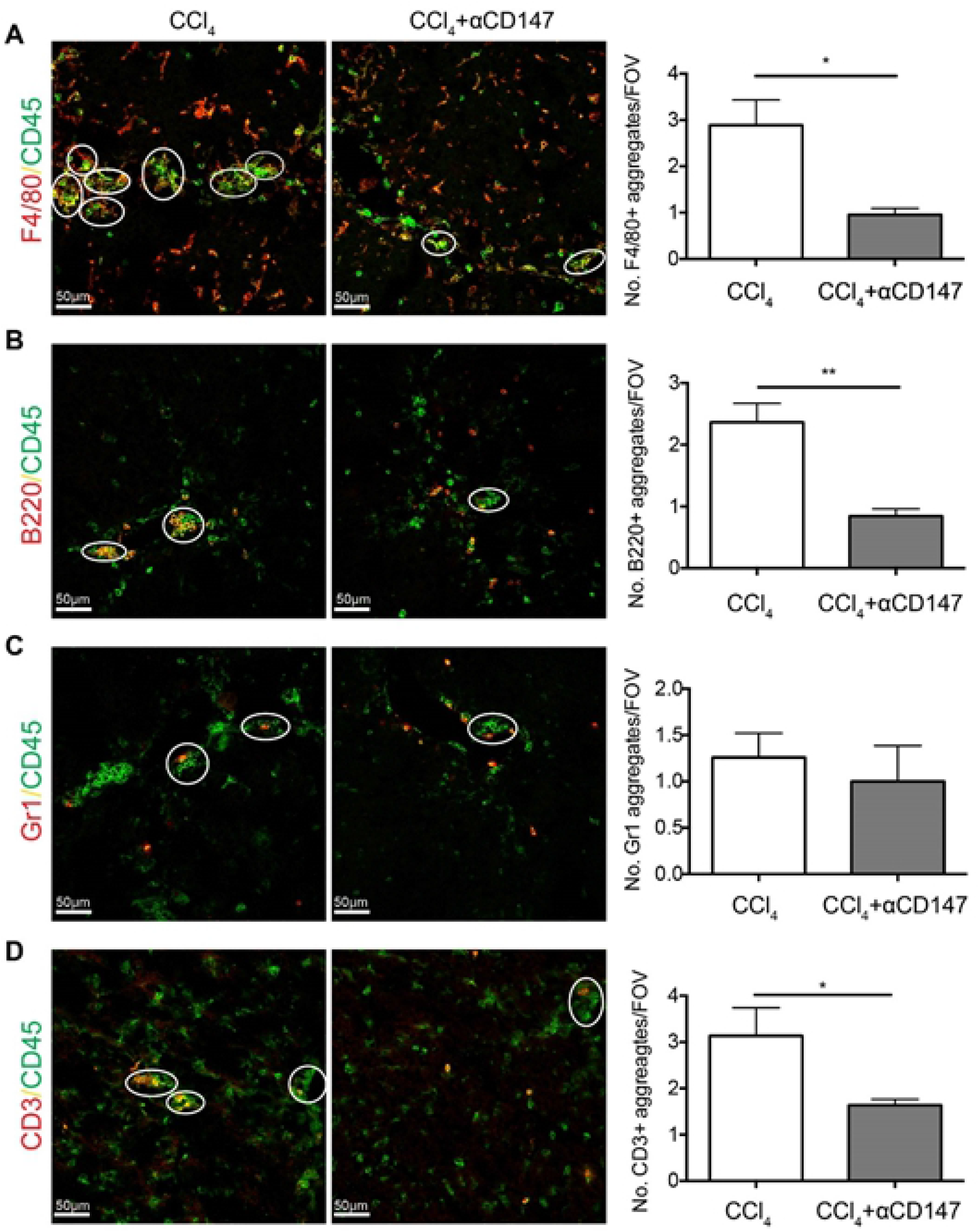
Anti-CD147 intervention reduces aggregation of macrophages, B cells, T cells and granulocytes during liver injury. CCl_4_ treated C57Bl/6 mice liver tissue was stained for CD45 colocalisation with (A) F4/80, (B) B220, (C) Gr1 or (D) CD3 to identify immune subset specific aggregates. Representative images (left panel) and quantified immune cell specific aggregates (right panel; calculated as at least one cell of interest per aggregate) in mice treated with CCl_4_ alone or CCl_4_ in conjunction with anti-CD147 intervention. CCl_4_ = injury alone, αCD147 = anti-CD147 in CCl_4_ injury.

## Discussion

This study has demonstrated that with progressive inflammation-associated tissue injury, immune cells cluster and contribute directly to the magnitude of the injury. Our novel discovery is that with liver injury CD147 is increased on the surface of leukocytes and then mediates cell-cell aggregation that determines the extent of liver injury. However, if we blocked CD147 with a mAb then CD45+ cell numbers in the liver remain unchanged but leukocytes are no longer found in aggregates. Importantly, we have already reported there is a significant reduction in liver injury seen with anti-CD147 mAb^(15)^. Further, CD147 mediated leukocyte aggregation appears to cause or significantly exacerbate injury as aggregate formation proceeds the development of significant fibrosis^(15)^. Therefore, this is not just a reduction in aggregation and inflammatory markers (AST/ALT) but also a reduction in resultant fibrosis.

All intrahepatic leukocyte subpopulations (CD4^+^, CD8^+^, NK, B-cell and macrophages) rapidly increase CD147 surface protein and total mRNA expression with liver injury. Therefore, this data shows that following liver injury, circulating and intrahepatic leukocytes increase CD147 expression. Subsequently, it has been shown leukocytes undergo activation^(57)^, aggregations are formed and then contribute to injury. It is unknown if leukocyte activation occours concurrent or prior to aggregation. The recruitment of inflammatory cells from the periphery to sites of liver injury is important in the pathogenesis of liver injury^(57)^ but its relationship to aggregation and dependence on CD147 expression was unrecognised. Based on these results CD147 is likely pivotal the development of inflammation in liver injury partly through the previously unrecognized role in leukocyte aggregate formation.

In liver injury anti-CD147 intervention leads to a reduction in immune cell aggregation characterised by a reduction in serum transaminases and total tissue MMP activity^(15)^. Importantly, there is no significant change in the total number of CD45+ cells infiltrating the liver. This suggests that the cells are not proliferating to form clusters but rather they aggregate with injury. Further, we have shown that the clusters that diminish with anti-CD147 intervention in injury contain B-cells (B220/CD19) and macrophages/Kupffer cells (F480). Importantly, the isotype antibody controls have the same phenotype as wild-type animals. Importantly, we have shown these leukocyte clusters form in progressive human liver injury irrespective cause.

CD147 has a number of binding interactions. The CD147 binding partners; α3β1, α6β1, CD44, LFA-1, ICAM-1, sydnedcan-1 and hyaluronan all have known roles in immune cell retention at sites of inflammation. The functional significance of CD147 protein binding interactions in the aggregation phenotype is unknown, likely to be pivotal and clearly needs to be determined. Multiple proteins are known membrane-binding partners of CD147 (see Background)^(29, 30, 38–40)^. Further, many of the CD147 complex proteins are found on leukocytes and/or have known roles in leukocyte retention at sites of inflammation (including CD98^(38)^, β1-integrin^(38)^, CD43^(58)^, LFA-1^(28, 43, 58)^, ICAM-1^(43)^, sydnecan-1^(28, 58)^, and hyaluronan^(39, 59))^. Additionally, CD147 through interactions with CD98^(38, 39)^ and/or β1-integrin^(38)^ is known to mediate homotypic cell clustering of leukocytes. Therefore, CD98^(38, 39)^, β1-integrin^(38)^, LFA-1^(28, 43, 58)^ and ICAM-1^(43)^ are the most promising protein binding interactions to study that are likely involved in the aggregation phenotype.

Therefore, we have identified CD147 as a potential therapeutic target to minimize exacerbation of liver injury via reduction of leukocyte aggregate accumulations in the liver tissue. Further work is required to determine the role of CD147 in liver fibrogenesis resultant from persistent inflammatory insult.

---

**Supplementary Figure 1.**
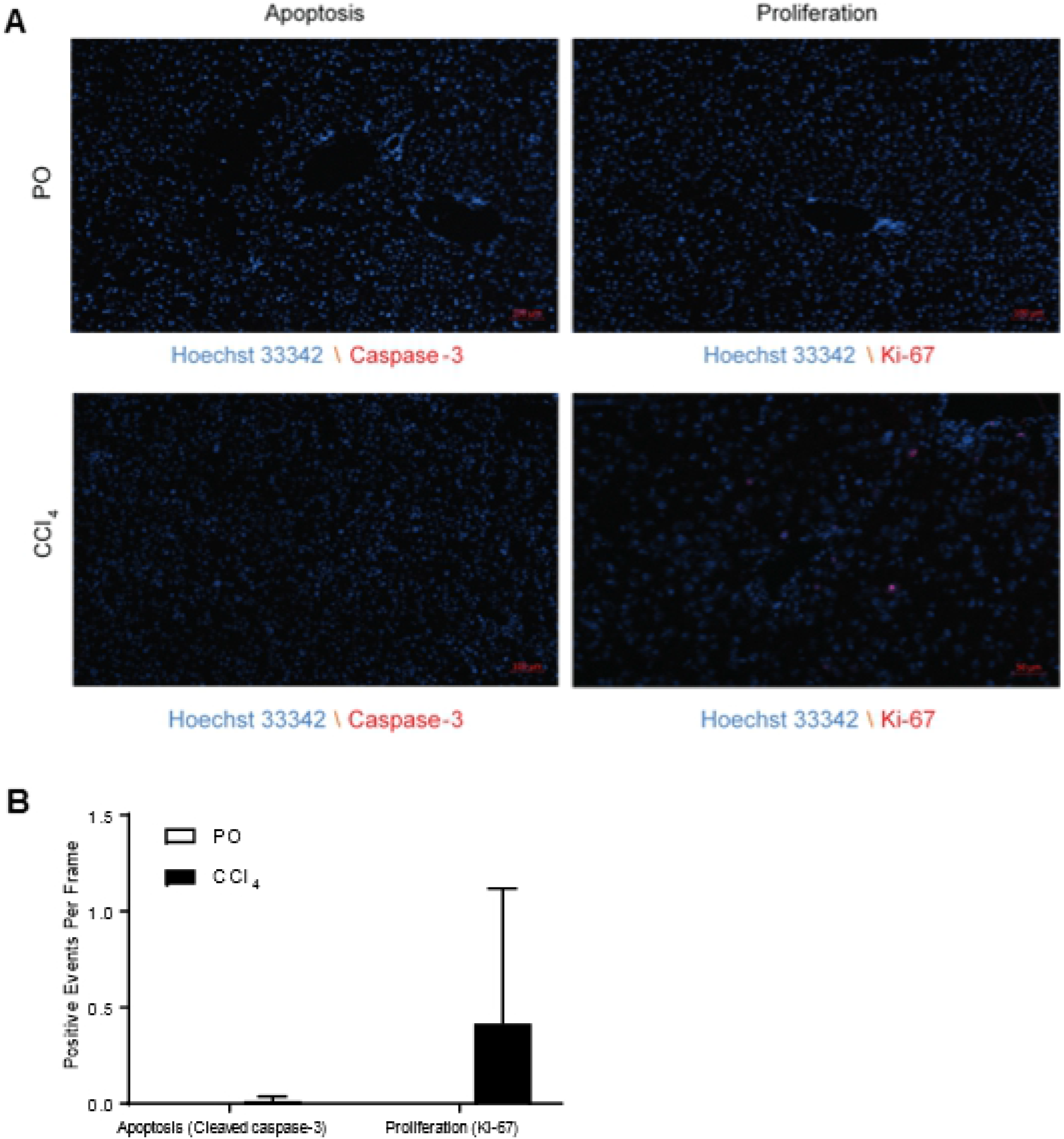
Apoptosis and proliferation within immune cell aggregates. In CCl_4_ treated C57Bl/6 mice and healthy controls, liver tissue was stained with cleaved caspase-3 and Ki-67 to assess apoptosis and proliferation within aggregates, respectively. (A) The number of positive events per frame where bar graphs represent mean ± SEM. (B) Representative images of apoptosis (left panels) and proliferation (right panels) from healthy control mice (top panels) and CCl_4_ treated mice (bottom panels). Mann-Whitney test was performed to assess significance from control.

